# AAV vectors with enhancer-controlled ITR promoters

**DOI:** 10.64898/2025.12.03.692098

**Authors:** Yanjiang Zheng, Luzi Yang, Kaiyu Zhou, Qirun Wang, Tiange Li, Jiahao Wu, jiangwei Qin, Fei Gao, Pingzhu Zhou, Tian Lu, Yangpo Cao, Yimin Hua, Yuxuan Guo, Yifei Li

**Affiliations:** Key Laboratory of Birth Defects and Related Diseases of Women and Children of MOE, Department of Pediatrics, West China Second University Hospital, Sichuan University, Chengdu, Sichuan, 610041, China; Institute of Cardiovascular Sciences, School of Basic Medical Sciences, Peking University Health Science Center, Beijing, China; State Key Laboratory of Vascular Homeostasis and Remodeling, Peking University, Beijing, China; Pulmonary Vascular Diseases Center, Beijing Anzhen Hospital, Capital Medical University, Beijing 101118, China; Department of Cardiovascular Surgery, West China Hospital, Sichuan University, Chengdu, Sichuan 610041, China; Institute for Cardiovascular Health, School of Medicine, Shanghai University, Shanghai 200444, China; Department of Biochemistry and Molecular Biology, School of Basic Medical Sciences, Peking University Health Science Center, Beijing, China; Department of Pharmacology, Joint Laboratory of Guangdong-Hong Kong Universities for Vascular Homeostasis and Diseases, School of Medicine, Southern University of Science and Technology, Shenzhen, Guangdong, 518033, China

**Keywords:** Adeno-associated virus, inverted terminal repeats, Enhancer

## Abstract

Adeno-associated virus (AAV) is limited by its packaging capacity and the unwanted promoter activity of inverted terminal repeats (ITRs). Here we utilized mini-enhancers to regulate ITR promoters and drive AAV cargo expression in the absence of the canonical promoters, which not only solved the ITR issue but also released more payload for the cargo. As an example, this new design successfully enabled robust, tissue-specific SpCas9 gene editing via a single AAV, which otherwise requires a dual-AAV system due to the large size of SpCas9.

## Main

Recombinant adeno-associated virus (rAAV) is a leading gene delivery vector in in vivo gene therapy. The payload of rAAV is limited to ∼4.7 kbp, which is the DNA length between the two inverted terminal repeats (ITRs). When AAV is used to deliver transgenes beyond its payload, the transgenes could be split into two pieces for a dual-AAV system, which raises efficacy and safety concerns due to the increased AAV dosage^1^. Alternatively, major transgene components could be carefully engineered to reduce their sizes, although with no guarantee of success^2^.

Conventionally, AAV transgenes are composed of a promoter, a coding sequence (CDS) and a 3’ untranslated region (3’UTR) with the polyadenylation signal (pA)^3^. Despite many efforts to reduce the size of each component, the minimal essential elements and maximal effective payload of a single AAV particle remain unclear. ITRs are essential for vector packaging, cell transduction and DNA replication^4^. In addition, ITRs are reported to initiate low-level transcription in an RNA polymerase II (Pol II)-dependent manner^5,6^. This ITR promoter activity is usually considered as an unwanted adverse factor, which could be blocked by incorporating additional pA or Woodchuck hepatitis virus posttranscriptional regulatory elements (WPREs) that further reduce the effective payload of AAV^7,8^. Alternative approaches that simultaneously solve the issues of packaging capacity and ITR activity remain untested.

Enhancers are cis-regulatory elements that determine the strength and spatiotemporal features of promoters^9^. Enhancers could be engineered to less than 100 bp while maintaining their activities in a position- and direction-independent manner, which was extensively studied in massively parallel reporter assays (MPRA) ^9,10^ and artificial-intelligence-based enhancer design^11^. However, whether enhancers could modulate ITR promoters and drive AAV transgene expression in the absence of conventional promoters remain to be determined.

As a means to test this hypothesis, a promoter-free AAV vector was constructed to express Cre recombinase directly downstream of the ITR (Fig. 1a). The cytomegalovirus (CMV) enhancer, a classic ubiquitously active enhancer^12^, was incorporated downstream of the 3’UTR and PolyA (Fig. 1a). These vectors were packaged into AAV9 and 5e10 vector genome (vg) AAV vectors were subcutaneously injected into postnatal day 1 (P1) mice harboring a loxp-stop-loxp-tdTomato reporter^13^. Seven days later, tdTomato signals were observed in ∼40% cells in the liver treated with AAV lacking the enhancer, validating the hepatic tropism of AAV9 and the basal promoter activity of ITR (Fig. 1b-c). Strikingly, the CMV enhancer profoundly enhanced the Cre reporter activity not only in the liver but also in the heart, lung, spleen and kidney (Fig. 1b-c). The directionality and distance of CMV enhancer relative to the ITR-Cre transgene exerted little impact on this effect (Fig. 1a-c), which is characteristic of an enhancer element.

**Fig.1.**
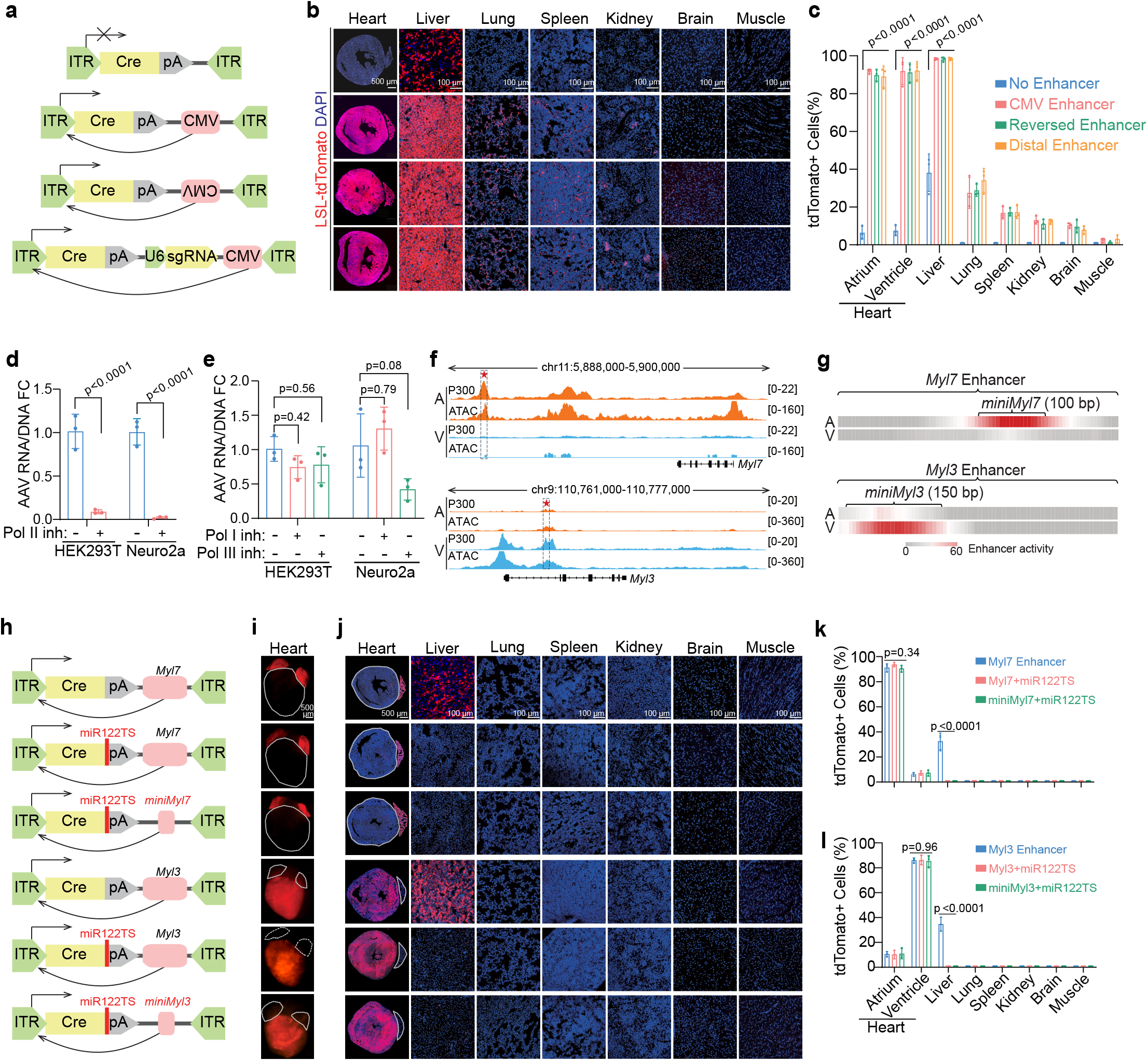
The ITR promoter can be modulated by mini-enhancers to regulate AAV transgene expression. **a**, Schematic of the ITR promoter-based AAV constructs. Cre CDS was positioned directly downstream of ITR without other canonical promoters. The CMV enhancer was placed in either forward or reverse orientation at varying distances relative to the ITR. **b-c**, Images (**b**) and quantifications (**c**) of tdTomato-positive cells in the indicated organs. LSL-tdTomato reporter mice were subcutaneously injected with AAV9 (5e10 vg) at P1 and analyzed a week later. N = 3 animals per group. **d-e**, RT-qPCR analysis of ITR promoter activity in HEK293T and Neuro2a cells upon RNA polymerase (Pol I∼Pol III) inhibitor treatment. Cells were treated with AAV-DJ packaging the ITR-Cre vectors with an MOI of 10^4^ vg/cell. 48 hours after AAV treatment, the cells were treated with 50 μM α-amanitin (Pol II inhibitor, **d**), 10 μM CX-5461(Pol I inhibitor, **e**) or 20 μM ML-60218 (Pol III inhibitor, **e**) before analysis. N = 3 cell culture repeats per group. **f**, Integrative genomic views of the murine *Myl7* and *Myl3* genes with P300 ChIP-seq and ATAC-seq signals in the heart. Star and box indicate the chamber-specific enhancers. **g**, MPRA of 5 bp-deletion libraries of mutant enhancers in the heart. **h**, Schematic of the AAV constructs incorporating the miniMyl7 or miniMyl3 enhancers and the liver-detargeting miR122TS in 3’UTR. **i-j**, Images of whole hearts (**i**) or sections of indicated organs (**j**). Dashed lines delineate the boundaries of heart chambers. **k-l**, Quantification of tdTomato-positive cells in the indicated organs. N = 3 animals per group. Data in **c, d, e, k, l** are presented as mean ± standard deviation. One-way ANOVA in **c, k, l**. Unpaired student’s t-test in **d-e**. ITR, inverted terminal repeat. CMV, cytomegalovirus. pA, polyadenylation signal. CDS, coding sequence. LSL, loxp-STOP-loxp. A, atrium. V, ventricle. Inh, inhibitor. FC, fold change. ATAC, assay for transposase-accessible chromatin. MPRA, massively parallel reporter assay. miR122TS, microRNA-122 target sequence. 3’UTR, 3’ untranslated region.

Next, 5e11 vg AAV9 vectors were tail-vein injected into 6∼10-week-old adult mice. Seven days later, CMV-enhancer-induced tdTomato signals were validated in the liver, heart, lung, spleen and kidney (Supplementary Fig. 1a-b). The vectors were also packaged into the AAV-DJ^14^, which was used to infect HEK293T or Neuro2a cells. Real-time quantitative PCR (RT-qPCR) validated Cre mRNA expression in these cell lines, which were suppressed by the Pol II inhibitor α-amanitin^15^, but not Pol I or Pol III inhibitors (Fig. 1d-e). Therefore enhancer-activated ITR promoters could drive AAV transgene expression independent of animal ages, AAV serotypes, administration routes or cell types.

Because the ITR-Cre vector was robustly expressed in the heart (Fig. 1b-c), this study next focused on the heart to test tissue-specific enhancers. Integrative analysis of the assay for transposase-accessible chromatin using sequencing and P300 chromatin immunoprecipitation sequencing previously revealed *Myl7* and *Myl3* enhancers in mice that were preferentially active in the cardiac atria or ventricles, respectively (Fig. 1f)^10^. AAV-based MPRAs using systemically mutagenized enhancer libraries identified core activity regions in *Myl7* and *Myl3* enhancers of ∼100 bp or ∼150 bp (Fig. 1g)^10^, which were termed miniMyl7 and miniMyl3 enhancers. *Myl7* and *Myl3* enhancers, and their miniature derivatives, specifically activated the ITR-Cre vectors in atrium and ventricles, respectively (Fig. 1h-l). The AAV transduction efficiency (0∼95% cells) in the designated chambers were titratable by adjusting AAV dosages (Supplementary Fig. 1c-d). No Cre reporter signals were detected in the lung, spleen, kidney, brain, or skeletal muscle (Fig. 1j-l). The leaky activity in the liver was eliminated by incorporating the microRNA-122 target sequences (miR122TS) in 3’UTR (Fig. 1h-l)^13,16^. Therefore, mini-enhancer-regulated ITR promoters could achieve tissue-specific AAV expression using only ∼100 bp payload.

Next, *Streptococcus pyogenes* Cas9 (SpCas9)-mediated cardiac gene editing was performed to showcase the application of enhancer-regulated AAV-ITR vectors. SpCas9 is the prototypic, heavily optimized, Cas9 nuclease that is limited by its large size^17^. The CDS of SpCas9 (4,107 bp) together with a U6-sgRNA gene (351 bp) occupies 95% AAV payload, leaving only ∼300 bp for the promoter and 3’UTR and thereby making it particularly challenging to be delivered by tissue-specific all-in-one AAV vectors. To solve this problem, here an AAV vector was built to express SpCas9 by the miniMyl7 enhancer (100 bp) together with a miR122TS-containing 3’UTR (217 bp) in the absence of a canonical promoter (Fig. 2a). 1e11 vg ITR-SpCas9-miniMyl7 vectors were injected into P1 mice for tissue collection at P30. By amplicon sequencing analysis^18^, three sgRNAs targeting the coding regions of *Scn5a*, a core gene regulating cardiac electrophysiology ^19,20^, lead to ∼20% insertion and deletion (indel) rates in the atria while less than 10% in the ventricles (Fig. 2b). The indel rates depended on AAV dosage (Fig. 2c) in the atria, while less than 1% indel rates were detected in the liver, lung, spleen, kidney, brain or skeletal muscles in the high-dose group (Fig. 2b, d). Immunofluorescence and western blot analysis verified SCN5A protein depletion predominantly in the atrium (Fig. 2e), further justifying tissue specificity.

**Fig 2.**
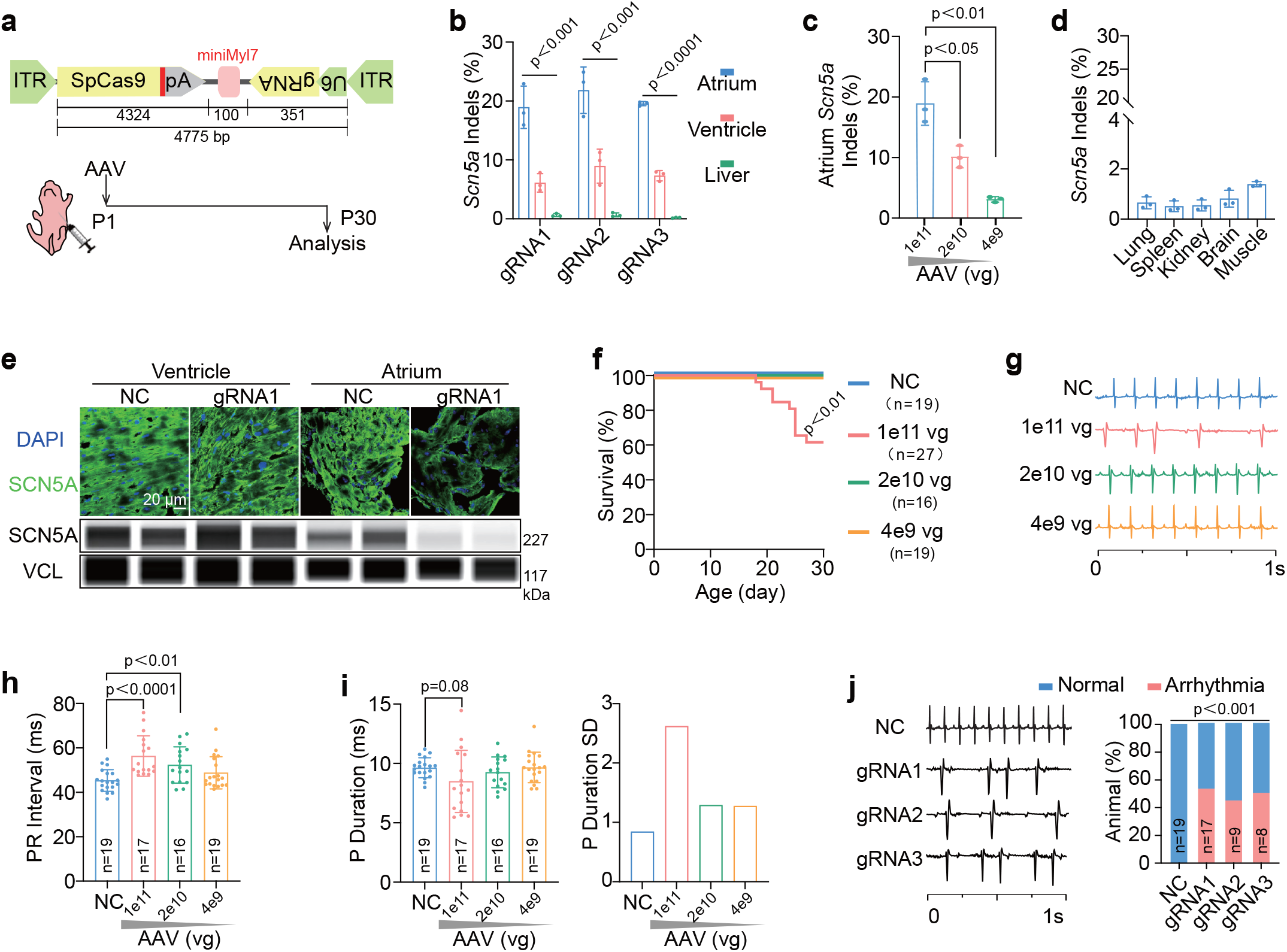
The ITR promoter enhances effective AAV payloads for single-AAV delivered SpCas9-mediated gene editing specifically in the cardiac atrium. **a**, Schematic of the all-in-one AAV vector expressing SpCas9 by the ITR promoter and the miniMyl7 enhancer and the experimental design of animal validation. **b-d**, Amplicon sequencing analysis of gene editing rates in the sgRNA targeted loci in the indicated organs or AAV doses. Three doses of ITR-SpCas9-miniMyl7 vectors were injected into P1 mice before tissue analysis at P30. **b** and **d** are data with 1e11 vg AAV. N=3 animals per group. **e**, Immunofluorescence (up) and western blot (down) analysis of SCN5A protein in heart chambers. **f**, Survival curve of mice with indicated AAV doses. N=16∼27 animals per group. **g**, Representative ECG traces of AAV-injected mice. **h-i**, Quantification of ECG traces. N = 16∼19 animals per group. **j**, ECG traces and quantification of animals treated with each of the three *Scn5a*-targeting sgRNAs. N = 8∼19 animals per group. Data in **b, c, d, h, i** are presented as mean ± standard deviation (SD). In **b**, one-way ANOVA. In **c, h, i**, Unpaired student’s t-test. In **f**, log-rank test. In **j**, Fisher’s exact test. SpCas9, *Streptococcus pyogenes* Cas9. Indel, Insertion and deletion. NC, negative control. VCL, vinculin. ECG, electrocardiogram. PR interval, time between the P wave and the following R wave in ECG.

Animals underwent sudden cardiac death starting at 2-3 weeks after AAV injection in a dose-dependent manner (Fig. 2f). Electrocardiogram detected arrhythmic decoupling between the P waves (atrial electrical signals) and QRS complexes (ventricular electrical signals) while the PR intervals were prolonged in the high-dose group (Fig. 2g-h). The mean of P wave durations decreased while their standard deviations increased (Fig. 2i). AAVs carrying each of the three sgRNAs caused similar electrophysiological phenotypes (Fig. 2j). Immunohistochemistry detected no obvious changes in Masson trichrome staining and Hematoxylin & Eosin staining of both cardiac chambers (Supplementary Fig. 2a). Echocardiogram detected no phenotypes in cardiac contractility and morphology (Supplementary Fig. 2b-c). Together, the single-AAV-delivered SpCas9-mediated *Scn5a* knockout in atria successfully recapitulated the expected phenotypes of atrial arrhythmia and atrioventricular blocks.

In summary, this study demonstrated that mini-enhancer-regulated ITR promoters could replace conventional Pol II-dependent promoters to drive AAV expression. This new design naturally solved the issue with unwanted ITR transcription while released more payloads for large cargos. With the presence of two ITRs, bicistronic vectors were expected to benefit most from this AAV design. With the accumulation of MPRA data and artificial-intelligence-assisted enhancer engineering, modular incorporation of mini-enhancers into the AAV-ITRs would likely empower future innovation of AAV gene therapy.

## Supporting information

Supplementary infromtion

## Methods

See supplementary information online for materials and methods.

## Data availability

Plasmids are available on Addgene. Sequencing data are available from the Sequencing Read Archive. Source data are provided with this paper.

## Contributions

Conceptualization, Y.Z., Y.G., Y.L., and T.L.;

Experimental studies, Y.Z., L.Y., K.Z., Q.W., T.L., J.W., T.L. and Q.W.;

Data collection and analysis, Y.Z., L.Y., Y.C., K.Z., T.L. and P.Z.;

Supervision, Y.L., Y.G., Y.C., Y.H. and F.G.;

Manuscript writing, Y.G. and Y.Z., with input from all authors.

## Competing interests

Enhancer-regulated ITR-promoter AAV vectors and their applications have patents applied by West China Second University Hospital and Peking University. All authors were provided with the full paper for comments and critiques before submission. The other authors declare no competing interests.

## Acknowledgements

This study was funded by the Noncommunicable Chronic Diseases—National Science and Technology Major Project (2025ZD0547100 to Y.L. and 2024ZD0526500 to Y.Z.), National Natural Science Foundation of China (82200265 to Y.Z., 82470251 to Y.Z., 82470247 to Y.C., 32400942 to Y.C.) and Beijing Natural Science Foundation (F252059/25FS1588 to Y.G. and F.G.).

## Notes

### Competing Interest Statement

The authors have declared no competing interest.

