## Supplementary infromtion for "AAV vectors with enhancer-controlled ITR promoters"

Zheng et al.

Supplementary Information

#### Supplementary Figure

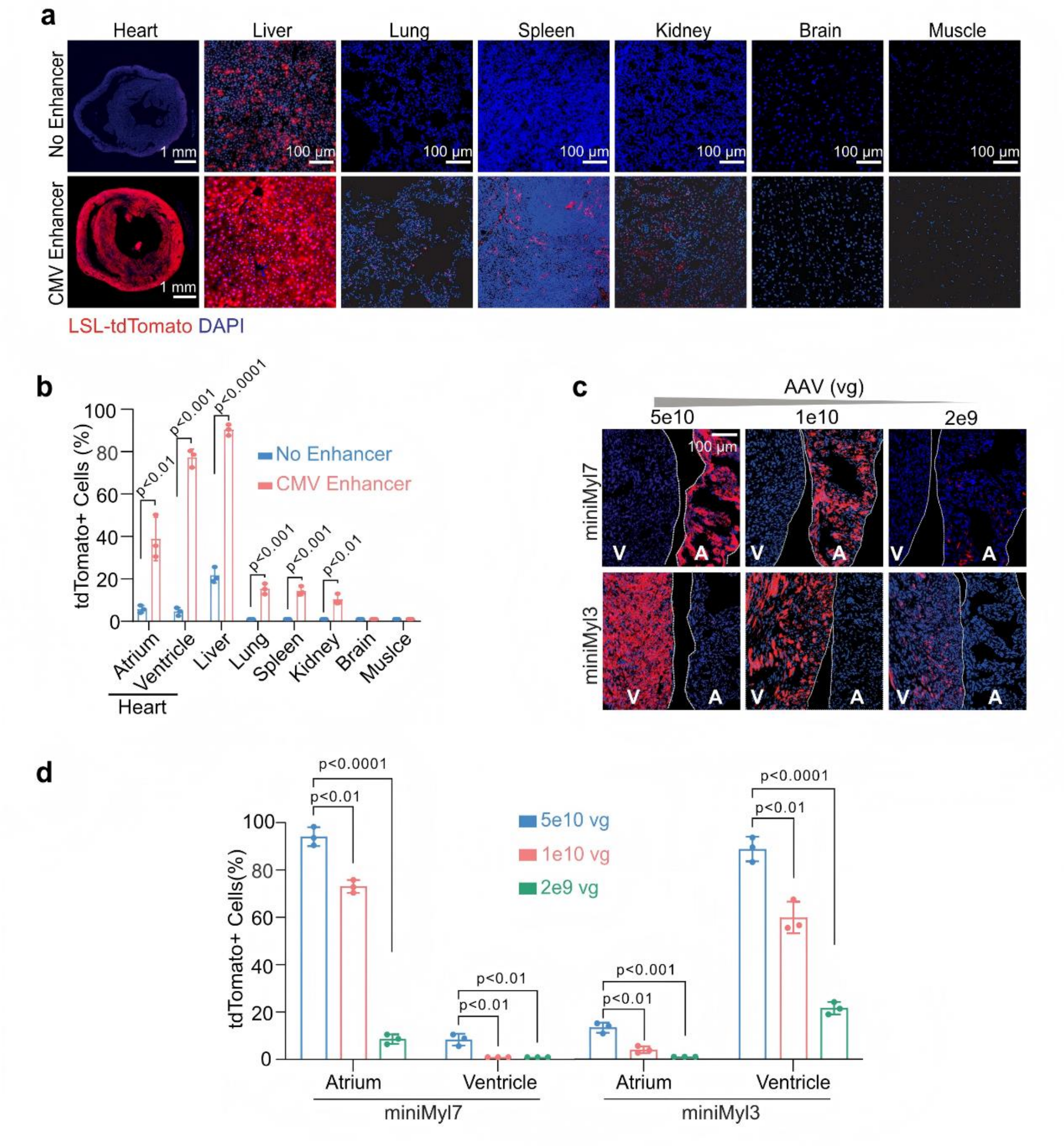

**Supplementary Fig.1: Enhancer-regulated ITR promoter drives AAV transgene expression in adult mice in a dose-dependent manner.**

**a-b,** Image (**a**) and quantification (**b**) of LSL-tdTomato activation in 6~10-week-old mice upon intravenously injection with 5e11 vg AAV9-ITR-Cre-CMV vectors. The indicated organs were analyzed at 7 days post-injection. **c-d,** Image (**c**) and quantification (**d**) of LSL-tdTomato activation in the heart upon injection with varying doses of ITR-Cre vectors harboring the miniMyl7 or miniMyl3 enhancers. Dashed lines delineate chamber borders. N = 3 animals per group. Data in **b, d** are presented as mean ± standard deviation. Unpaired Student’s t-test. vg, vector genome. LSL, loxp-stop-loxp. A, atrium. V, ventricle.

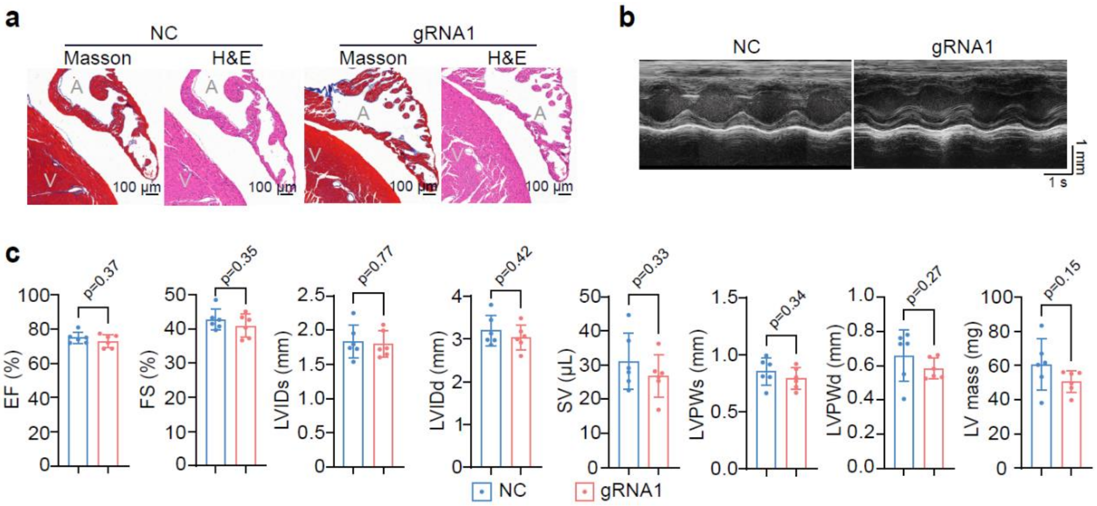

**Supplementary Fig.2: Single-AAV-delivered SpCas9-mediated *Scn5a* editing does not trigger overt cardiac structural or contractile defects in the heart.**
**a,** Histological analysis of Masson staining and Hematoxylin & Eosin (H&E) staining in the heart. **b-c,** M-mode echocardiogram images (**b**) and quantification (**c**) of the key cardiac contractility and morphology parameters. N = 6 animals per group. Mean ± standard deviation. Unpaired Student's t-test. NC, negative control. EF, ejection fraction. FS, fractional shortening. LVIDs, left ventricular internal dimension at end-systole. LVIDd, left ventricular internal dimension at end-diastole.

#### Supplementary Table

##### Supplementary Table 1: Full plasmid sequences

The list and full sequences of the AAV plasmids are provided in a separate file.

##### Supplementary Table 2: Nucleic acid sequences

| Application | Name | Nucleic acid sequence (5'->3') |
|---|---|---|
| Cre qPCR of cDNA | Primer- F | GGACATGTTCAGGGACAGGC |
| | Primer- R | TGGCTTGCAGGTACAGGAGG |
| human Gapdh qPCR of cDNA | Primer- F | GAGTCAACGGATTTGGTCGT |
| | Primer- R | GACAAGCTTCCCGTTCTCAG |
| mouse Gapdh qPCR of cDNA | Primer- F | TGACCACAGTCCATGCCATC |
| | Primer- R | TGACCACAGTCCATGCCATC |
| U6 qPCR of cDNA | Primer- F | CTCGCTTCGGCAGCACA |
| | Primer- R | AACGCTTCACGAATTTGCGT |
| mouse Tnni3 qPCR of gDNA | Primer- F | GCCAGGTTGACTGAAGAGACAG |
| | Primer- R | CTGGTTTGGAGAGGTTTATTCTGC |
| human Gapdh qPCR of gDNA | Primer- F | TTCCACCGCAAAATGGCCCCTC |
| | Primer- R | CCAGACACCCAATCCTCCCGGT |
| Scn5a-targeted SpCas9 editing | gRNA1 | CAGCTTCCGTAGGTTCACCC |
| | gRNA2 | GAAGGGGCTGAGGACGTACA |
| | gRNA3 | GAGGACGTACAAGGCATTGG |
| Amplicon sequencing | Scn5a-F1 | ACACTCTTTCCCTACACGACGCTCTTCCGATCTAGCTTCCTTCCAGGCAGCCT |
| | Scn5a-R1 | ACTGGAGTTCAGACGTGTGCTCTTCCGATCTCTCACGGCTCTCCTGTGAGG |
| | Scn5a-F2 | ACACTCTTTCCCTACACGACGCTCTTCCGATCTGACAGCCGTCTCCAGACCTA |
| | Scn5a-R2 | ACTGGAGTTCAGACGTGTGCTCTTCCGATCTCGTACCCCAGGACAGATGCA |

**Supplementary Table 3: Key reagents**

| Antibody | Use | Supplier | Cat. No. |
|---|---|---|---|
| [Rabbit]Anti-Scn5a | Western blot (1:50) Immunofluorescence (1:200) | Affinity Biosciences, USA | DF13217 |
| [Rabbit]Anti-Vinculin | Western blot (1:200) | Abcam, USA | EPR8185 |
| Anti-Rabbit IgG, AF-488 | Immunofluorescence (1:500) | ThermoFisher, USA | A-21206 |
| Anti-Rabbit IgG, HRP | Western blot (1:200) | ProteinSimple, USA | 042-206 |
| **Reagent** | **Use** | **Supplier** | **Cat. No.** |
| DAPI | Immunofluorescence (1:1000) | Abcam, USA | ab285390 |
| WGA-iFluor647 | Immunofluorescence (1:200) | AAT Bioquest, USA | 25559 |
| Isoflurane | Anesthesia | RWD, China | R510-22-10 |
| PBS | Cell culture | Gibco, USA | C20012500BT |
| DMEM | Cell culture | Gibco, USA | C11995500BT |
| Pen/Strep | Cell culture | Gibco, USA | 15140122 |
| FBS | Cell culture | Sigma, USA | F8313 |
| OptiMEM | Cell transfection | Gibco, USA | 31985070 |
| PEI | Cell transfection | Sigma, USA | 408727 |
| PEG-8000 | AAV production | Solarbio, China | P8260 |
| Benzonase | AAV production | Yeasen, China | 20157ES50 |
| CX-5461 | Cell culture (10 μM) | MedChemExpress, USA | HY-13323 |
| α-amanitin | Cell culture (50 μM) | MedChemExpress, | HY-19610 |
| ML-60218 | Cell culture (20 μM) | MedChemExpress, | HY-122122 |
| **Name** | **Species** | **Supplier** | **Cat. No.** |
| HEK293T | human | Meisen Cell, China | CTCC-DZ-0321 |
| Neuro2a | mouse | Meisen Cell | CTCC-003-0093 |

### Supplementary Methods

#### Mouse

Animal experiments were approved by the Institutional Animal Care and Use Committee of West China Second University Hospital of Sichuan University. LSL-tdTomato mice (Stock NO.: T002249) were obtained from GemPharmatech, China. C57BL/6 mice were obtained from Chengdu Dossy Experimental animals. Mice were maintained in individually ventilated cages under controlled environmental conditions, including a temperature of 23 ± 1℃, relative humidity of 50 ± 5%, and a 12-h dark/light cycle. Food and water were provided ad libitum.

The administration route of AAVs was determined by the age of the mice. Neonatal (P1) mice received subcutaneous injections, while adult mice were administered via tail vein injection. Anesthesia was induced with 3% isoflurane (R510-22-10, RWD). Mice under the age of 8 days were sacrificed by decapitation, whereas older animals were euthanized by cervical dislocation under anesthesia.

#### Plasmid

Plasmid names and full sequences are summarized in Supplementary Table 1. Plasmids for cell culture transfection and AAV packaging share the same backbone. The miR122TS (Addgene #117384)[1], Myl7 enhancer, and Myl3 enhancer DNA sequences were synthesized by Youkang Biotech, China. The sequences of CMV enhancer, reversed CMV enhancer, U6-gRNA, SpCas9 (Addgene #87115)[2], miniMyl7, and miniMyl3 were amplified by PCR.

The U6-gRNA1-U6-gRNA2-TnT-promoter element was deleted from the AAV-U6gRNA1-U6gRNA2-TnT-Cre plasmid (Addgene #87682)[3] to generate an AAV-ITR-Cre plasmid lacking conventional promoters. Subsequently, the CMV enhancer, reversed CMV enhancer, Myl7 enhancer, and Myl3 enhancer elements were individually ligated into the RsrII site of the AAV-ITR-Cre plasmid backbone to generate the ITR-Cre-CMV, ITR-Cre-reverseCMV, ITR-Cre-Myl7, and ITR-Cre-Myl3 plasmids. The U6-gRNA cassette was inserted into the ITR-Cre-CMV plasmid via the AscI restriction site to generate the ITR-Cre-U6-gRNA-CMV construct. The miR122TS element was cloned into the ITR-Cre-Myl7 and ITR-Cre-Myl3 plasmids using BamHI sites, resulting in the ITR-Cre-miR122TS-Myl7 and ITR-Cre-miR122TS-Myl3 plasmids. The Myl7 and Myl3 sequences were replaced with miniMyl7 and miniMyl3, respectively, to create the ITR-Cre-miR122TS-miniMyl7 and AAV-CremiR122TS-miniMyl3 plasmids.

Cre CDS in the ITR-Cre-miR122TS-miniMyl7 plasmid was replaced with the SpCas9 sequence to generate the all-in-one ITR-SpCas9-miR122TS-miniMyl7 plasmid. The *Scn5a* gRNA sequences were designed using the CRISPRpick tool (Broad Institute GPP Portal) and were listed in Supplementary Table 2. The U6-gRNA elements were commercially synthesized by Youkang and inserted into the ITR-SpCas9-miR122TS-miniMyl7 backbone via the RsrII site to produce the ITR-SpCas9-miR122TS-U6-gRNA-miniMyl7 plasmid.

**AAV package**

AAV was produced in house[3] or with Packgene Biotech, China. A total of 7 μg of AAV genome (target plasmid), 7 μg of either AAV9 or AAV-DJ Rep/Cap plasmid, and 20 μg of the pHGT1-adeno/dF6 helper plasmid were co-transfected into HEK293T cells cultured in one 15-cm dish. The transfection was performed using a mixture of 1.8 mL Opti-MEM (Sigma, USA, #408727) and 170 μL polyethylenimine (Sigma, #408727) transfection reagent. After 66 - 72 hours, the cells were harvested and resuspended in AAV lysis buffer (20 mM Tris pH 8.0, 1 mM $MgCl_2$, 150 mM NaCl). Cell lysis was achieved through three cycles of freezing at -80 °C and thawing at 37 °C. The crude lysate was then purified by ultracentrifugation through a discontinuous gradient of four OptiPrep densities using a T70ti rotor at 68,000 rpm for 150 minutes at 4 °C. The purified AAV was concentrated in PBS containing 0.001% Pluronic® F-68 (Thermo Fisher, USA, #24040032). Viral genome titer was determined by real-time quantitative PCR (RT-qPCR) using a standard curve generated from serial dilutions of the backbone plasmids.

**Tissue collection and fluorescence imaging**

Mice were euthanized with $CO_2$ or isoflurane, and then the tissues were excised. For whole-heart imaging, hearts were cleaned by cold PBS. Then the hearts were put on a glass-bottom dish, and immediately imaged by a upright fluorescence microscope (M165, Leica, Germany).

For cryo-sectioning, tissues were fixed overnight at 4 °C in 4% paraformaldehyde in PBS. The fixed tissues were then transferred to 15% sucrose for several hours, followed by immersion in 30% sucrose overnight at 4 °C. The tissues were next embedded in Tissue Freezing Medium (General Data, TFM-5) and frozen at -80 °C. Tissues sections were cut at a thickness of 10 μm using a cryostat (Thermo

Scientific, Microm HM550) for subsequent staining.

Cryo-sections were rinsed twice with PBS for fluorescence imaging. Relevant reagents were listed in Supplementary Table 3. For direct staining, samples were incubated with Wheat Germ Agglutinin (WGA, 1:200) and DAPI (1:1,000) in PBS for 1 hour at room temperature, followed by two PBS washes. For immunofluorescence staining, sections were permeabilized with PBST (0.1% Triton X-100 in PBS) for 10 minutes, blocked with 4% BSA in PBS for 10 minutes, and incubated overnight at 4°C with primary antibody against SCN5a (Affinity Biosciences, USA, #DF13217; 1:200). The next day, samples were rinsed three times with PBS and incubated for one hour at room temperature with secondary antibody (1:500) and DAPI (1:1,000). All samples underwent final PBS washes before being mounted with Diamond Antifade Mountant (Thermo Fisher, P36965) for imaging.

For cryosection fluorescence imaging, images were taken by an Olympus FV3000 inverted laser scanning confocal microscope with ×10, ×20, and ×60 objectives. Cell size and number were quantified using ImageJ software (version 1.54f).

**Cell culture**

HEK293T and Neuro2A cells were obtained from Meisen Cell, China. Cells were cultured in high glucose DMEM containing 10% FBS and 1×Penicillin-Streptomycin solution at 37°C and 5% CO2 with saturating humidity. These cells were plated at a density of $1.5\times10^5$ cells per well in 24-well plates at 16 hours before AAV transduction at the MOI of $10^4$ vg/cell. 48 hours after AAV treatment, cells were washed twice using PBS and then treated with Pol I (CX-5461,10 μM), Pol II (α-amanitin, 50 μM) or Pol III (ML-60218, 20 μM) inhibitors. The cells were collected 6-8 hours after RNA transcription inhibitor treatment before collecting RT-qPCR analysis.

**RT-qPCR analysis**

For RNA quantification, total RNA was extracted using the TransZol Up Plus RNA Kit (TransGen Biotech, China, ER501-01). 1 μg RNA was reverse transcribed to cDNA by HiScript III All-in-one RT SuperMix Perfect for qPCR (Vazyme, China, R333-01). Genomic DNA was removed from RNA before reverse transcription. For DNA quantification, genomic DNA (gDNA) was extracted with TIANamp Genomic DNA Kit (TIANGEN, China, GDP304-03). RT-qPCR was performed on a Aria Mix Real-Time PCR system (Agilent Technologies, USA) using 2x Taq Pro Universal SYBR qPCR Master Mix (Q712-Q2, Vazyme Biotech). In cell culture experiments, AAV-expressed

RNA levels were normalized to AAV DNA levels to assess ITR promoter activity. Primers for RT-qPCR were listed in Supplementary Table 2.

**Amplicon sequencing analysis**

To extract DNA for amplicon sequencing, tissues were digested overnight at 55 °C with 0.5 mL of DNA digestion buffer (50 mM Tris–HCl pH 8.0, 1 mM EDTA pH 8.0, 100 mM NaCl, 1% SDS) containing proteinase K. Then genomic DNA was precipitated by adding 1 mL of 100% isopropanol, collected by centrifugation, and washed with 70% ethanol. Then the DNA pellet was dissolved in 50 μL of water.

DNA samples were amplified by PCR using primers in Supplementary Table 2. Additional rounds of PCR were performed to attach next-generation sequencing adapters. The amplicons were purified by electrophoresis on a 2% agarose gel, followed by cleanup with a FastPure Gel DNA Extraction Mini Kit (Vazyme, DC301), and eluted in 25 μl of $H_2O$. Sequencing was carried out on the Illumina NovaSeq X Plus platform. The data was analyzed using CRISPResso2 (crispresso.pinellolab.org) to calculate indel rates[4].

**Western bolt**

Tissues were homogenized in 50 μl RIPA lysis buffer (Epizyme Biotech, China, PC101) supplemented with 1× protease inhibitor cocktail, followed by sonication for 90 seconds. The lysates were centrifuged at 12,000 × g for 15 minutes at 4 °C, and the supernatant was collected. Protein concentration was determined using a BCA Protein Assay Kit (Epizyme, ZJ102).

Western blot was performed by capillary electrophoresis (Jess, Proteinsimple, USA). For each sample, 10 μg protein was mixed with primary antibodies and secondary antibodies, then loaded into an 8 × 25 capillary cartridge. The cartridge was subsequently placed into the Wes automated protein detection system for analysis. Relevant reagents were listed in Supplementary Table 3.

**Histology analysis**

Heart tissue specimens were fixed in 10% neutral buffered formalin to preserve morphology, followed by routine processing through a graded series of ethanol for dehydration, xylene for clearing, and infiltration with molten paraffin wax. The processed tissues were then embedded in paraffin blocks, which were sectioned at a thickness of 3 μm using a rotary microtome. The resulting ribbons were floated on a water bath, mounted onto glass slides.

Masson's trichrome staining and H&E staining were performed by Wuhan Servicebio Technology, China. For Masson's trichrome staining, sections were processed using the Masson dye solution set (G1006, Servicebio) in accordance with the manufacturer's protocol. Briefly, sections were incubated in Masson A solution overnight and subsequently rinsed under tap water. A working Masson solution was prepared by mixing Masson B and Masson C at a 1:1 ratio. The sections were then stained in this solution for 1 min, differentiated in 1% hydrochloric acid alcohol for several seconds, and rinsed again with tap water. This was followed by sequential immersion in Masson D for 10 min, Masson E for 1 min, and Masson F for 30 s. After a final rinse with 1% glacial acetic acid, the sections were dehydrated through two changes of anhydrous ethanol. Finally, the slides were cleared in xylene for 5 min and mounted with neutral balsam.

H&E staining was carried out using the H&E HD kit (G1076, Servicebio) following the manufacturer's protocol. Briefly, sections were treated with HD constant staining pretreatment solution for 1 min, followed by staining in hematoxylin solution for 3–5 min. After rinsing under running tap water, the sections were differentiated in hematoxylin differentiation solution for 3–5 s, rinsed again, blued in hematoxylin bluing solution for 3–5 s, and rinsed. Subsequently, the sections were immersed in 95% ethanol for 1 min and counterstained with eosin for 15 s. Finally, the samples were dehydrated through three changes of anhydrous ethanol, two changes of normal butanol, and two changes of xylene, then mounted with neutral balsam.

The stained sections were digitally scanned using a Pannoramic MIDI microscope slide scanner and analyzed by SildeViewer software (3DHISTECH, Hungary).

**Echocardiography & electrocardiography**

Mice were anesthetized with 3% isoflurane, maintained asleep under anesthesia with 1–1.5% isoflurane, and analyzed by operators blinded to the experimental groups. Echocardiography was performed using the Vevo 3100 ultrasound system (VisualSonics, Canada) and measured using M-mode echocardiography. For electrocardiography, lead II electrocardiograms were recorded using PowerLab hardware and analyzed with LabChart software (ADInstruments, Australia).

**Statistical analysis**

Statistical analyses and data visualization were conducted using GraphPad Prism version 10 for Windows (www.graphpad.com). The normality of data distribution was assessed using the Shapiro–Wilk test. For data satisfying normality, comparisons between two groups were performed with Student's t-test. Comparison between three or more groups were performed using one-way ANOVA. Survival analysis was carried out using Kaplan–Meier curves with log-rank test. The incidence of arrhythmia among animals was evaluated using Fisher's exact test. Unless otherwise noted, data are presented as mean ± standard deviation in bar plots following confirmation of normal distribution, with raw data points on the graph.
